# Integrated Clinical and Omics Approach to Rare Diseases: Novel Genes and Oligogenic Inheritance in Holoprosencephaly

**DOI:** 10.1101/320127

**Authors:** Artem Kim, Clara Savary, Christèle Dubourg, Wilfrid Carré, Charlotte Mouden, Houda Hamdi-Rozé, Hélène Guyodo, Jerome Le Douce, Laurent Pasquier, Elisabeth Flori, Marie Gonzales, Claire Bénéteau, Odile Boute, Tania Attié-Bitach, Joelle Roume, Louise Goujon, Linda Akloul, Erwan Watrin, Valérie Dupé, Sylvie Odent, Marie de Tayrac, Véronique David

## Abstract

**Purpose:** Holoprosencephaly (HPE) is a pathology of forebrain development characterized by high phenotypic and locus heterogeneity. Seventeen genes are known so far in HPE but the understanding of its genetic architecture remains to be refined. Here, we investigated the oligogenic nature of HPE resulting from accumulation of variants in different relevant genes.

**Methods:** Exome data from 29 patients diagnosed with HPE and 51 relatives from 26 unrelated families were analyzed. Standard variant classification approach was improved with a gene prioritization strategy based on clinical ontologies and gene co-expression networks. Clinical phenotyping and exploration of cross-species similarities were further performed on a family-by-family basis.

**Results:** We identified 232 rare deleterious variants in HPE patients representing 180 genes significantly associated with key pathways of forebrain development including Sonic Hedgehog (SHH) and Primary Cilia. Oligogenic events were observed in 10 families and involved novel HPE genes including recurrently mutated genes (*FAT1, NDST1, COL2A1* and *SCUBE2*) and genes implicated in cilia function.

**Conclusions:** This study reports novel HPE-relevant genes and reveals the existence of oligogenic cases resulting from several mutations in SHH-related genes. It also underlines that integrating clinical phenotyping in genetic studies will improve the identification of causal variants in rare disorders.

## Introduction

Holoprosencephaly (HPE1, OMIM #236100) is a severe developmental defect resulting from incomplete forebrain cleavage. The disease is characterized by incomplete separation of cerebral hemispheres with several anatomical classes ranging from microforms to alobar HPE.^1^ Affected individuals present with typical craniofacial midline defects of varying severity including proboscis, cleft lip and palate, ocular hypotelorism and solitary median incisor. HPE occurs in about 1 in 10,000 to 20,000 live births worldwide.^1^

The genetic basis of HPE remains unclear and different transmission patterns have been described including autosomal dominant, recessive and digenic inheritance.^1–3^ Most mutations associated with HPE display incomplete penetrance and variable expressivity, *i.e*. close relatives carrying the same pathogenic variant can be asymptomatic or present distinct HPE-spectrum anomalies.^1,4^ Mutations in the *SHH, ZIC2, GLI2* and *SIX3* genes are the most frequently reported ones in HPE and collectively account for 16 % of the cases. Pathogenic variants in *FGF8, FGFR1, DISP1, DLL1* and *SUFU* were found in 8 % of HPE cases.^5,6^ The other HPE genes reported so far are *TDGF1, FOXH1, TGIF1, CDON, NODAL, GAS1* and *STIL*, whose frequency is not established due to the small number of observed cases.^1–3^

Clinical genetic testing of HPE has improved, but approximately 70 % of familial cases remain without a clear molecular diagnosis. Most of known HPE genes belong to the Sonic Hedgehog (SHH) pathway, which represents the primary pathway implicated in the disease.^1,4,5,7^ Therefore, defective SHH-related processes are likely to be substantially involved in HPE.

Whole-exome sequencing (WES) has been very successful for Mendelian disease-gene discovery and differential diagnosis.^8^ WES analysis employs filtering approaches for candidate variant prioritization combined with comprehensive clinical evaluation. A variety of additional strategies has been developed to further improve the performance of WES in clinical settings. Collaborative platforms such as Matchmaker Exchange^9^ are used to search for recurrence in patients affected by similar phenotypes. Integrative variant-prioritization algorithms such as the Exomiser suite^10^ combine WES with different phenotype-driven approaches (based on clinical data and cross-species phenotype comparisons) and analysis of protein interactome data. As useful as they are, these strategies are limited: collaborative platforms are not efficient in case of very rare genetic diseases while pipelines such as Exomiser are not designed to study non-Mendelian disorders. Studying HPE faces these two challenges: (i) HPE live-born infants are excessively rare, and, (ii) HPE is characterized as a complex non-Mendelian trait, *i.e*. that cannot be explained by the presence of a single mutation.

Recent studies have highlighted that non-Mendelian disease phenotypes could present an oligogenic etiology and result from accumulation of inherited low-penetrance variants in multiple genes.^11^ However, such events are likely overlooked in clinical genetic studies if variants are inherited from a clinically unaffected parent.

In this study, we address the additional yield that can be obtained for HPE patients who underwent medical WES evaluation that failed to establish a molecular diagnosis. Given the wide clinical spectrum of the disease, as well as incomplete penetrance and variable expressivity of HPE mutations, we raised the possibility that the low diagnostic yield is partly due to the complex etiology of HPE and hypothesized that a part of unsolved HPE cases results from oligogenic events, *i.e*. accumulation of several rare hypomorphic variants in distinct, functionally connected genes.

Our study involved patients for whom no disease etiology could be determined by conventional diagnostic approaches. Similarly to previous WES studies^12,13^, we used clinically-driven prioritization approach to identify genes associated with specific clinical features as reported in gene-phenotype reference databases and mouse models. Complementarily, we developed and used a prioritization strategy based on gene co-expression networks of the developing human brain to select genes with spatio-temporal expression patterns compatible with those of known HPE genes. Finally, we used in-depth clinical phenotyping together with cross-species similarities to further strengthen the evidence of causality.

This study highlights novel HPE genes and identifies new disease-related pathways including the primary cilia pathway. Our findings also illustrate the high degree of oligogenicity of HPE and suggest that the disease requires a joint effect of multiple hypomorphic mutations.

## Materials & Methods

### Patient selection

Study protocol was approved by the Ethics Committee of Rennes Hospital. Patients diagnosed with HPE and relatives were recruited using the clinical database of Holoprosencephaly Reference Center of Rennes Hospital. Study participation involved informed written consent, availability of clinical data, and either DNA or peripheral blood sample. All patients were scanned for rare damaging mutations by targeted HPE gene-panel sequencing^5^ and for copy number variants (CNVs) using CGH-array and MLPA.

### WES and variant classification procedure

As part of routine diagnosis procedure to identify the genetic causes of HPE, patients and relatives underwent WES using standard procedures as previously described.^2,3^ WES data analysis and filtering protocols are described in the *Supplementary Appendix*. Our scheme for variant classification followed the American College of Medical Genetics and Genomics-Association (ACMG) guidelines.^14^ Patients for whom no genetic etiology had been identified in clinical settings were included in the study.

### Clinically-driven approach

We established two clinician-generated lists of relevant phenotypes reminiscent of HPE in human and mouse models respectively (*Table S3*). Genes associated with the phenotypes of interest were identified with publicly available clinical resources and associated ontologies. Human gene-phenotype associations were extracted from relevant databases (*Figure S1*) using R package *VarFromPDB* (https://github.com/cran/VarfromPDB). The Mouse Genome Informatics (MGI)^15^ database and a homemade workflow were used to retrieve genes associated with any of the corresponding phenotypes in mouse mutants. Human and mouse results were combined and redundancy was removed to establish a list of clinically-driven candidate genes associated with HPE-related anomalies.

### Identification of HPE-related genes by weighted-gene co-expression network analysis

We used Weighted Gene Co-expression Network Analysis (WGCNA)^16^ on the RNA-Seq data from the Human Development Biology Resource (HDBR)^17^ to identify genes sharing highly similar expression patterns with 4 classical genes associated to HPE *– SHH, SIX3, ZIC2* and *TGIF1 –* during cerebral development. Data from samples corresponding to forebrain, cerebral cortex, diencephalon, telencephalon, and, temporal lobe structures taken between the 4th and 10th post-conception weeks (pcw) were selected (*Figure S9*). RNA-seq data were analyzed with the iRAP pipeline (https://github.com/nunofonseca/irap). We used R package WGCNA to construct co-expression networks and identify modules of co-expressed genes. The detailed protocols for WGCNA analysis are described in the *Supplementary Appendix*. The Topological Overlap Matrix (TOM) matrix was used to establish a list of transcriptome-driven candidate genes sharing highly similar expression profiles with *SHH, ZIC2, SIX3* and *TGIF1*.

### Integration and in-depth analyses of results

The two prioritization schemes were combined with the WES results to identify a restricted list of rare variations located in disease-relevant genes (Figure 1). To determine significantly enriched biological processes and pathways, functional annotation was performed by *g:profiler* (http://biit.cs.ut.ee/gprofiler) and Bonferroni adjusted p-value were considered significant below 0.05 value (KEGG, REACTOME and Gene Ontology Biological Processes).

**Figure 1.**
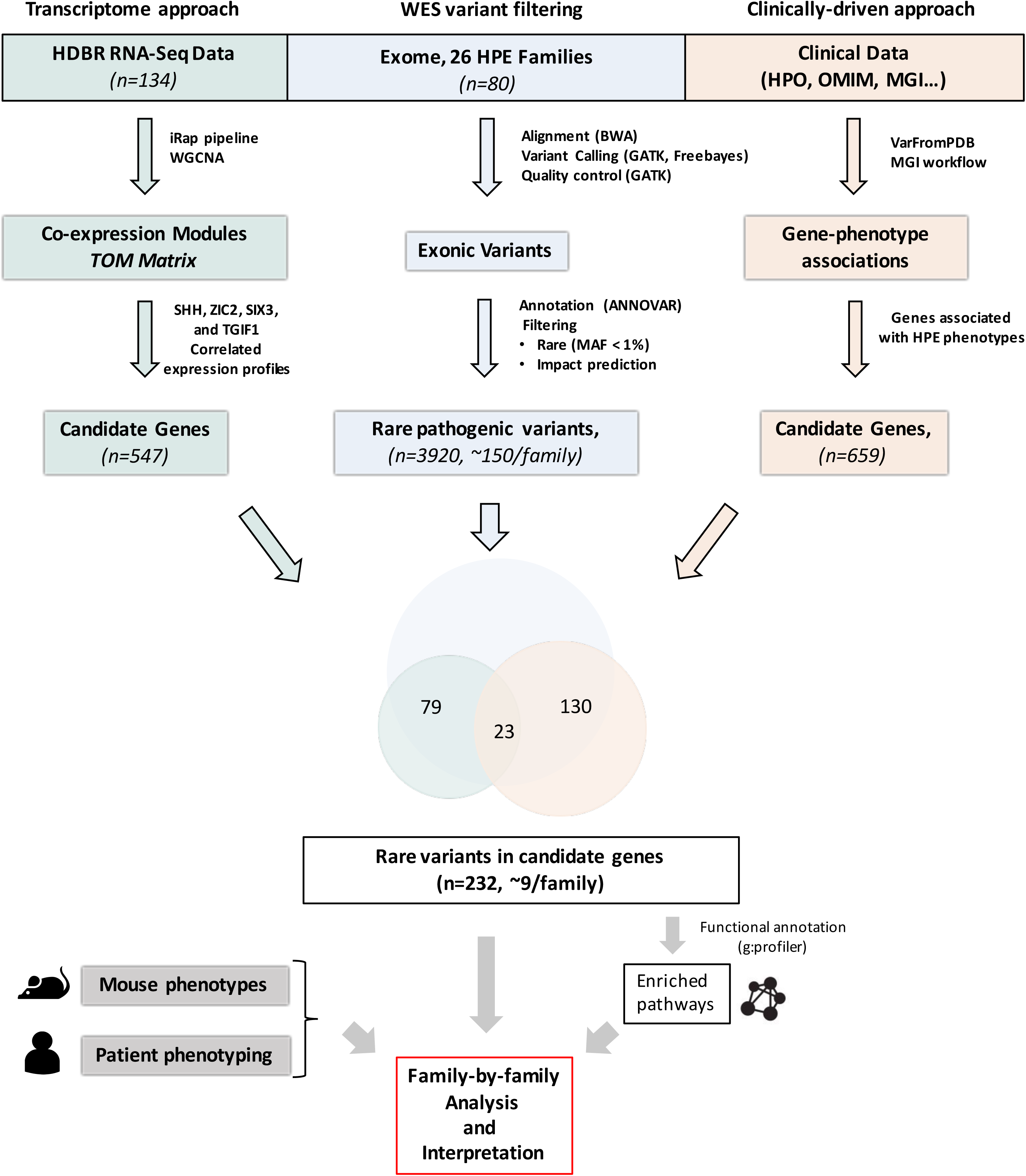
Flow chart illustrating the prioritization strategy. Classical WES analysis was performed (blue) and combined with two prioritization approaches: (1) based on gene co-expression networks (green) and (2) based on clinical knowledge (salmon). Details of the pipeline are also provided in the Supplementary Appendix. Variant overlaps were selected and further analysed by functional annotation analysis and on a family-by-family basis, by integrating a comprehensive clinical phenotyping of patients and exploration of cross-species similarities.

Further analyses of the candidate variants were performed on a family-by-family basis. Oligogenic events were defined as combinations of variants unique to the affected proband(s). Variants could be either inherited from the parents— at least one each from the mother and the father - or occur *de novo*. We explored common types of variant-based evidence for classifying pathogenic variants^18^ (*e.g*. population, computational, functional and segregation data). To further evaluate the impact of candidate genes, we performed deep clinical phenotyping to characterize similarities between unrelated patients and/or published knockout mice. Special attention was given to genes harboring distinct rare variants in at least two affected patients with striking phenotypic overlap. Phenotypic overlaps between patients and mouse mutants deficient for the corresponding candidate genes were also examined. The most interesting combinations of candidate variants in the affected probands were finally discussed during multidisciplinary meetings.

## Results

### Clinical findings

We assembled a cohort of 26 families representing a total of 80 individuals including 29 affected probands diagnosed with lobar (*n=3*), semilobar (*n=11*), alobar (*n=13*) or microform HPE (*n=2*) (Table 1). Common HPE clinical manifestations were observed among the probands and included cleft lip and palate (38 %), hypotelorism (34 %), microcephaly (31 %) and arhinencephaly (31 %). Ancestry analysis identified that 24 families were of European descent and two of South East Asia and African descent (*Figure S10*). 8 parents presented minor signs of midline facial anomalies and 3 parents were diagnosed with HPE microforms. Point mutations in known HPE genes were found in 13 families and a full heterozygous deletion *of SIX3* gene was detected in one family (*Figure S8*). As all anomalies of known HPE genes were inherited from asymptomatic or mildly affected parents, they were insufficient to explain the etiology of HPE in the corresponding families, indicating the presence of potential additional mutational events required for the disease to occur.

**Table 1.**
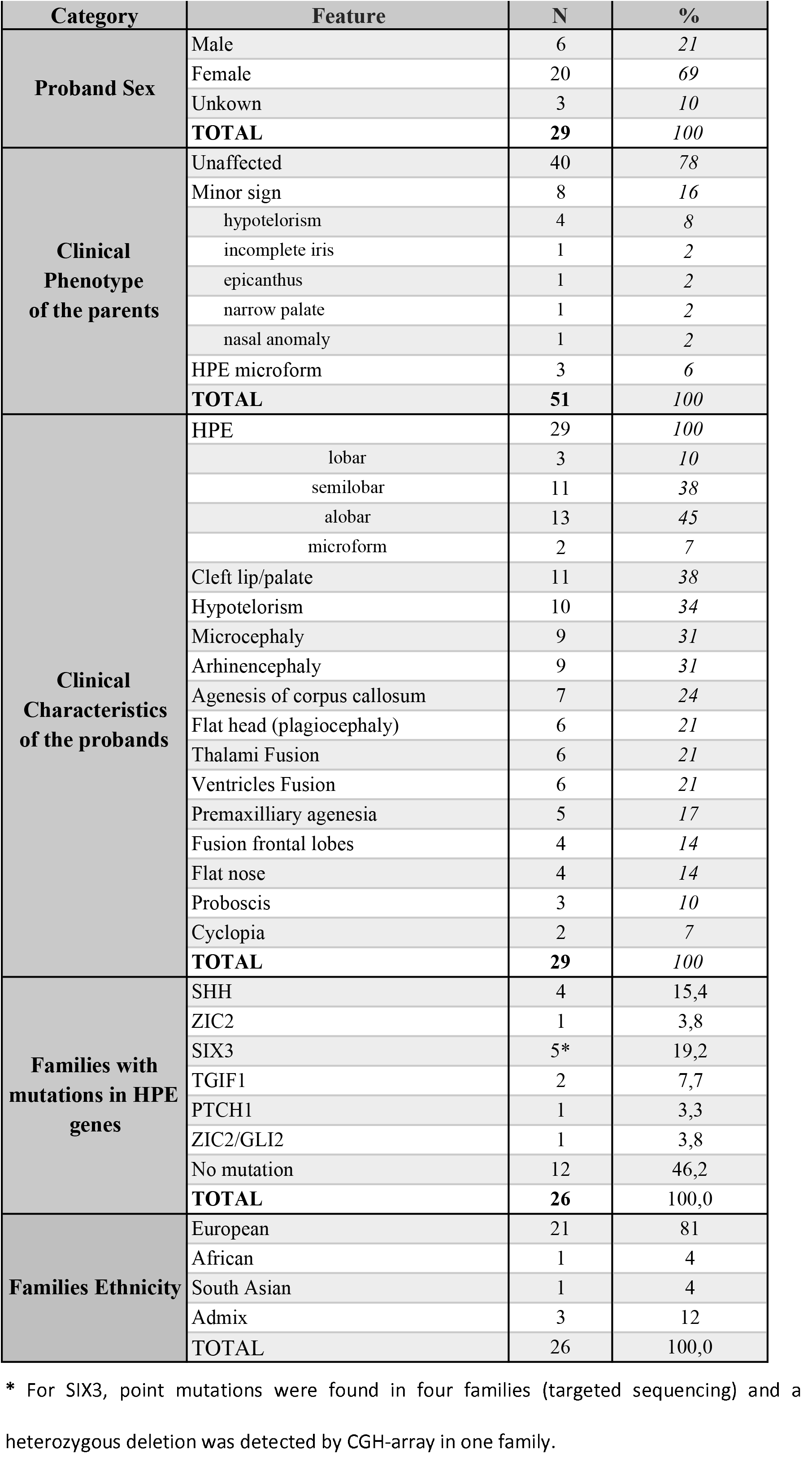
Clinical description of 26 HPE families.

| Category | Feature | N | % |
| --- | --- | --- | --- |
| Proband Sex | Male | 6 | 21 |
|  | Female | 20 | 69 |
|  | Unkown | 3 | 10 |
|  | **TOTAL** | **29** | **100** |
| Clinical Phenotype of the parents | Unaffected | 40 | 78 |
|  | Minor sign | 8 | 16 |
|  | hypotelorism | 4 | 8 |
|  | incomplete iris | 1 | 2 |
|  | epicanthus | 1 | 2 |
|  | narrow palate | 1 | 2 |
|  | nasal anomaly | 1 | 2 |
|  | HPE microform | 3 | 6 |
|  | **TOTAL** | **51** | **100** |
| Clinical Characteristics of the probands | HPE | 29 | 100 |
|  | lobar | 3 | 10 |
|  | semilobar | 11 | 38 |
|  | alobar | 13 | 45 |
|  | microform | 2 | 7 |
|  | Cleft lip/palate | 11 | 38 |
|  | Hypotelorism | 10 | 34 |
|  | Microcephaly | 9 | 31 |
|  | Arhinencephaly | 9 | 31 |
|  | Agenesis of corpus callosum | 7 | 24 |
|  | Flat head (plagiocephaly) | 6 | 21 |
|  | Thalami Fusion | 6 | 21 |
|  | Ventricles Fusion | 6 | 21 |
|  | Premaxillary agenesis | 5 | 17 |
|  | Fusion frontal lobes | 4 | 14 |
|  | Flat nose | 4 | 14 |
|  | Proboscis | 3 | 10 |
|  | Cyclopia | 2 | 7 |
|  | **TOTAL** | **29** | **100** |
| Families with mutations in HPE genes | SHH | 4 | 15,4 |
|  | ZIC2 | 1 | 3,8 |
|  | SIX3 | 5* | 19,2 |
|  | TGIF1 | 2 | 7,7 |
|  | PTCH1 | 1 | 3,3 |
|  | ZIC2/GLI2 | 1 | 3,8 |
|  | No mutation | 12 | 46,2 |
|  | **TOTAL** | **26** | **100,0** |
| Families Ethnicity | European | 21 | 81 |
|  | African | 1 | 4 |
|  | South Asian | 1 | 4 |
|  | Admix | 3 | 12 |
|  | **TOTAL** | **26** | **100,0** |

### HPE variants overview and identification of disease-related pathways

Combined clinically- and transcriptome-driven analysis of the exome data identified a total of 232 rare candidate variants in 180 genes (Figure 1 and Table S4). All variants presented a Minor Allele Frequency (MAF) below 1 % and were predicted to be highly deleterious to protein function (see *Supplementary*). 153 variants concerned genes associated with HPE phenotypes among which 32 were located in genes known to induce HPE-like phenotypes in mutant mice (*Table S5*). 102 variants were located in genes sharing expression profiles highly similar to those of HPE genes. Overlap between phenotype and gene co-expression network analysis contains 23 variants including 14 previously described mutations in known HPE genes (*SHH, ZIC2, SIX3, GLI2, TGIF1* and *PTCH1*).

Consistent with known disease etiology, functional profiling of the 180 genes revealed a significant enrichment for biological processes implicated in forebrain development (Table S6) including Sonic Hedgehog signalling pathway (*REAC:5358351, p-value = 2,79E^−5^; KEGG:04340, p-value = 1E^−4^*), Primary Cilia (*REAC:5617833; p-value = 1E^−6^; G0:0060271, p-value = 2E^−6^*) and Wnt/Planar Cell Polarity (PCP) signalling pathway (GO:0016055, p-value = 2E^−5^). The SHH pathway is the primary pathway implicated in HPE and the primary cilium is required for the transduction of SHH signalling^19,20^ while components of Wnt/PCP pathway regulate both SHH signalling and primary cilia.^19,21^

In-depth analyses highlighted 10 families with oligogenic events (Figure 2) clustered among 19 genes (Table 2) that functionally relate to disease-relevant pathways (Figure 3). These combinations of variants were unique to the affected probands. Main findings are presented below and full reports are available in the *Supplementary Appendix*.

**Table 2.**
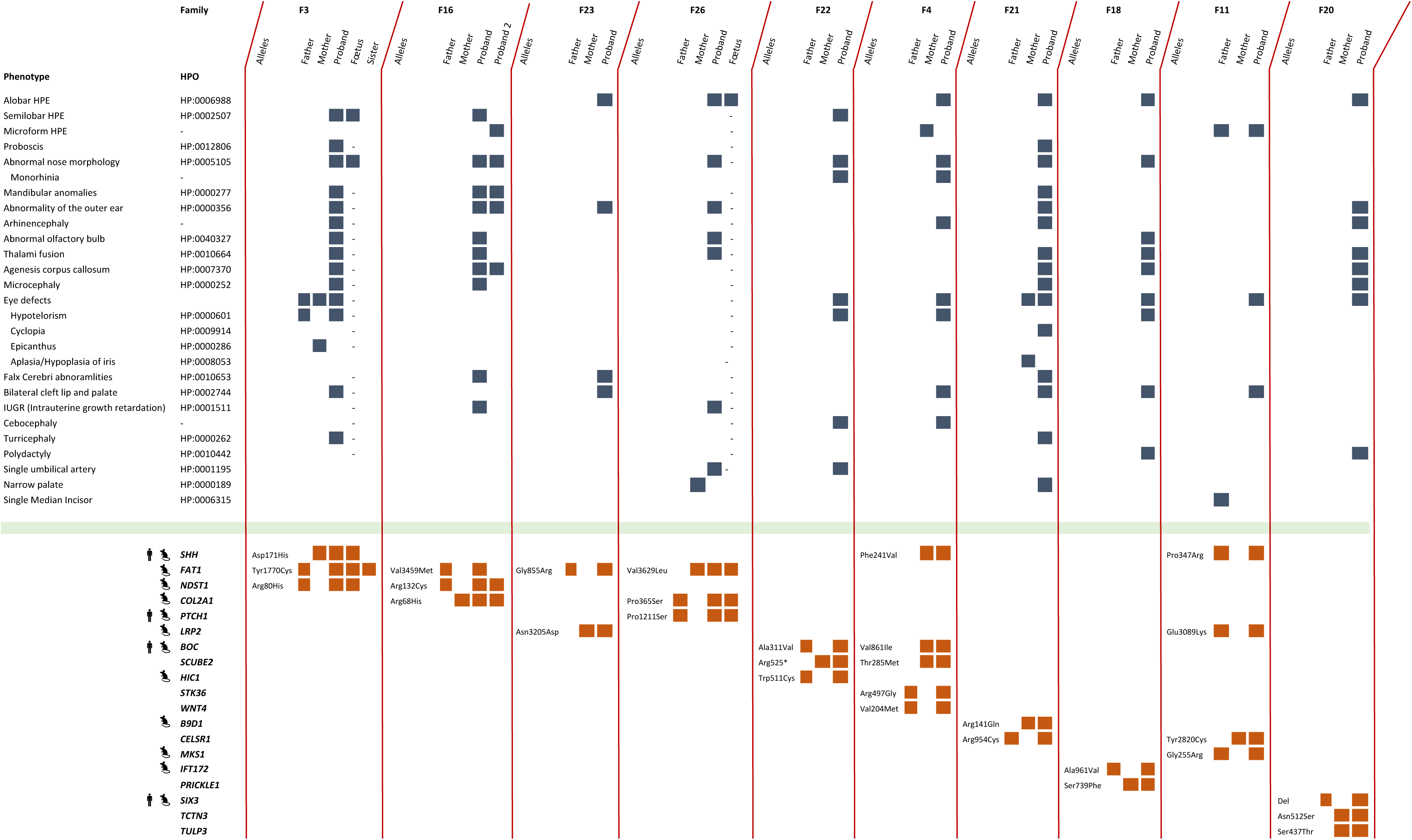
Comparison of clinical features in the studied families. Occurrences of phenotypes are marked with blue squares for each individual. Unrelated families are separated by red lines. Hyphen is used when no observation was possible on foetuses. Heterozygous variants in the different genes are marked with orange squares. Human symbols indicate that the genes are known HPE disease genes. Mouse symbols indicate the existence of mouse mutant exhibiting HPE for the corresponding gene.

**Figure 2.**
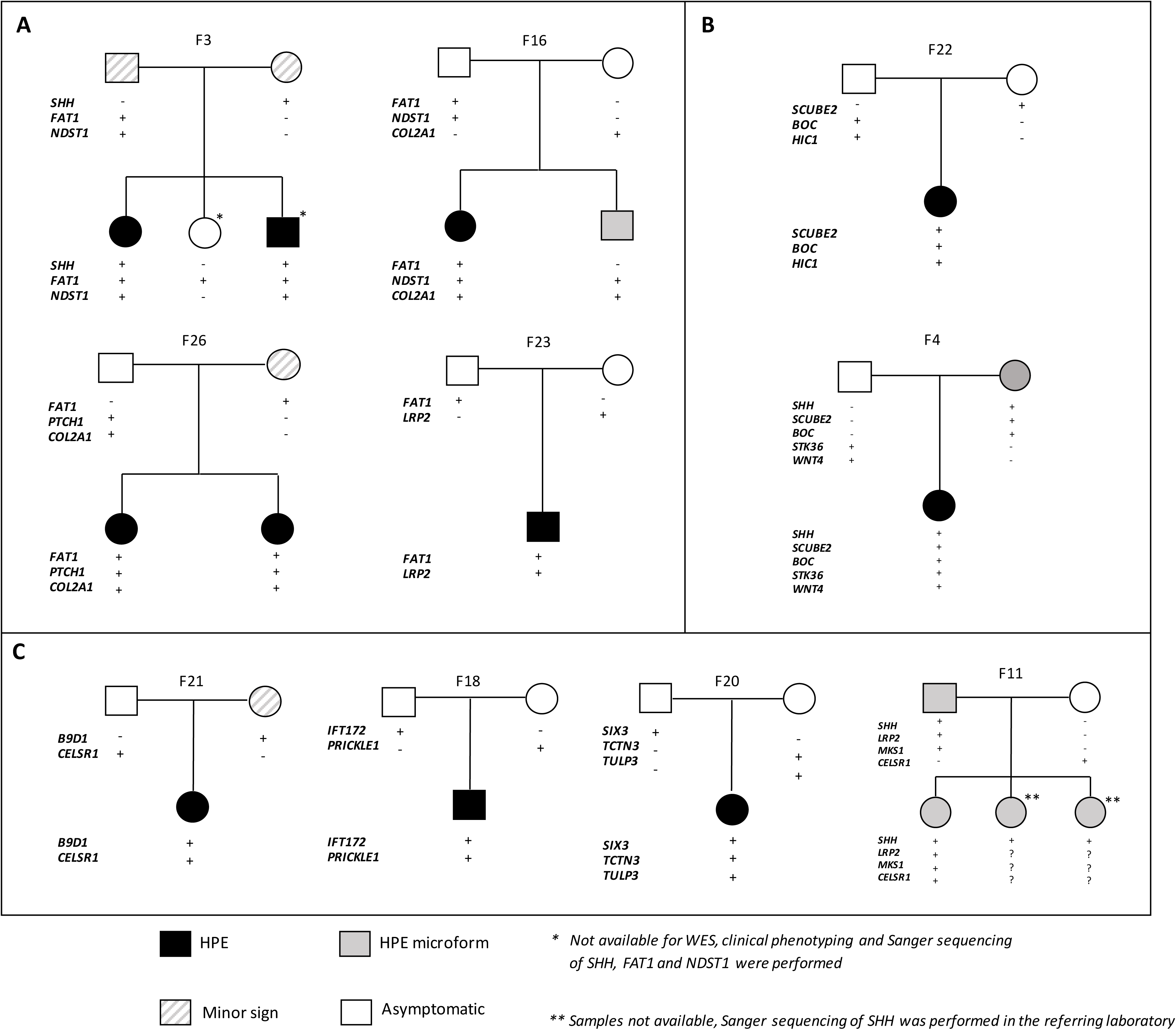
Oligogenic events reported in this study. Candidate genes are listed for each family. Individuals carrying/not carrying the variants are identified respectively by the *+/−*. Variants information is available in Table 2 and Table S4 of the Supplementary Appendix. **(A)** Oligogenic events involving *FAT1*. **(B)** Oligogenic events involving variants in *SCUBE2* and *BOC*.(***C***) Oligogenic events involving mutations in genes related to the primary cilium.

### Recurrent oligogenic events involving FAT1

4 different families, *i.e*. 15 % of the 26 families studied here, presented oligogenic events involving *FAT1* in combination with rare variants in known HPE genes (*SHH, PTCH1*), as well as in *NDST1, COL2A1* and *LRP2* genes (Figure 2A). FAT1 is a protocadherin and its knockdown in mouse causes severe midline defects including HPE,^22^ when in Drosophilia it has been shown to regulate the PCP pathway.^23^ *LRP2, NDST1* and *COL2A1* are all functionally relevant to the SHH pathway (Figure 3)*: NDST1* and *COL2A1* mice mutants exhibit HPE phenotype and reduced SHH signalling in the forebrain,^24,25^ while *LRP2* acts as an auxiliary receptor of SHH during forebrain development and its inactivation in mouse similarly leads to HPE phenotype.^26^

**Figure 3.**
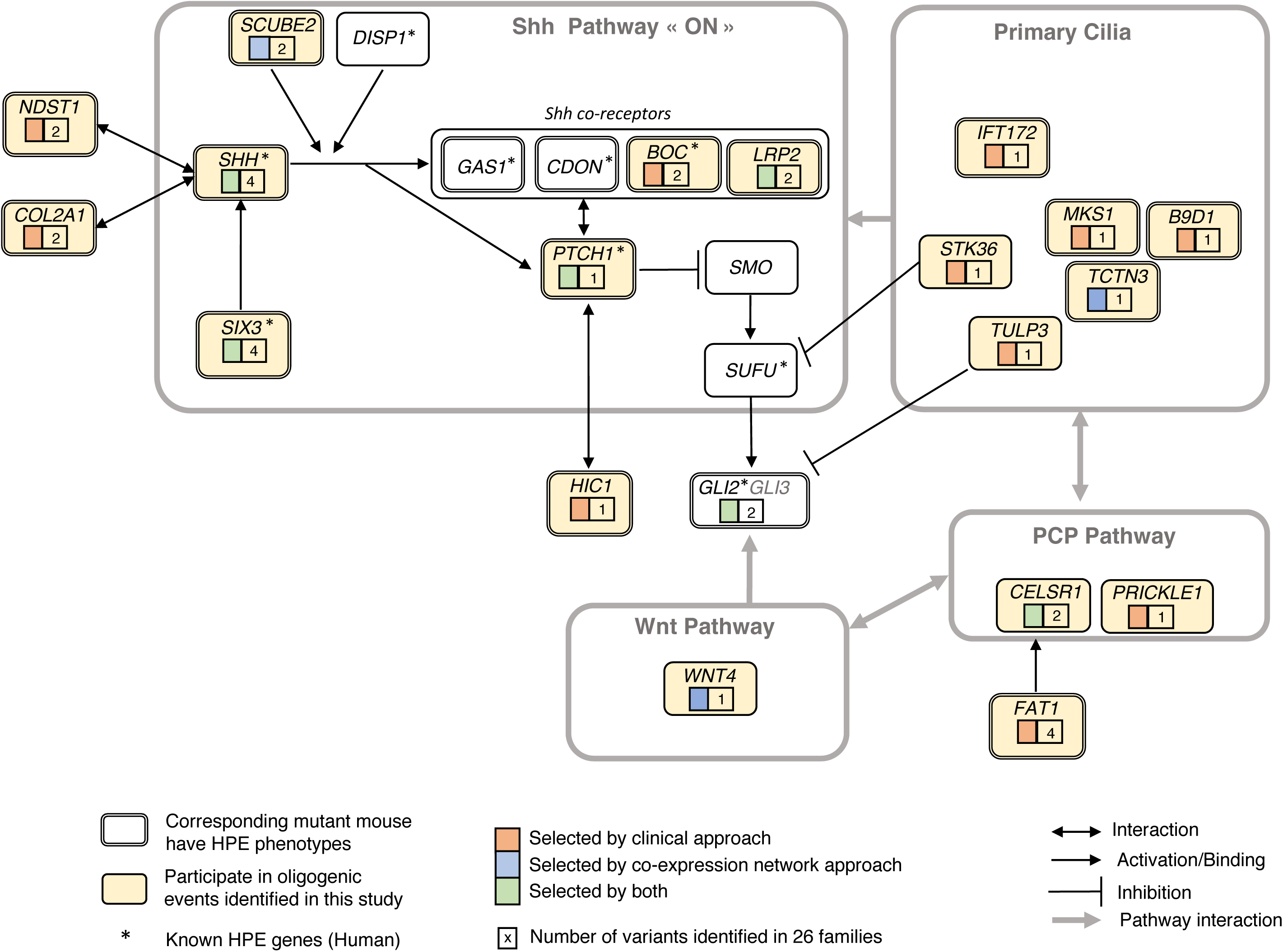
Implication of the candidate genes in the signaling pathways involved in HPE. Key affected pathways and genes are presented. Under each gene name, the selection methods (clinical or co-expression networks approach or both) is shown in the left panel and the number of variants for each gene is shown in the right panel. Genes known in HPE are marked with an asterisk, and genes for which corresponding mutant mouse have HPE phenotypes are surrounded by a double line. The genes implicated in an oligogenic events in the study are presented in yellow background.

Oligogenic events involved the following combinations: *SHH/FAT1/NDST1* (family F3), *FAT1/NDST1/COL2A1* (Family F16), *FAT1/COL2A1/PTCH1* (family F26) and *FAT1/LRP2* (Family F23) (Figure 2A and Table 2). The details are provided in the *Supplementary appendix, Case report* (*1*).

In family F3, Sanger sequencing of additional family members revealed that the *SHH/FAT1/NDST1* combination was unique to the affected individuals (Figure 2A). For family F16, only the foetus carrying the *FAT1/NDST1/COL2A1* combination was affected by semilobar HPE, while the sibling carrying *NDST1/COL2A1* variants presented only a microform (Figure 2A). These observations are fully consistent with the oligogenic inheritance model where accumulation of multiple variants in genes associated to HPE phenotypes and/or HPE-related molecular pathways is required.

### Recurrent oligogenic events involving SCUBE2/BOC implicated in SHH signalling

Similarly, 2 families presented oligogenic events implicating combined variants in *BOC* and *SCUBE2* genes (Figure 2B and Table 2). BOC is an auxiliary receptor of SHH and was recently reported as HPE modifier in Human.^27^ *SCUBE2* shares a highly similar expression pattern with *SHH* and *SIX3* and is implicated in the release of *SHH* from the secreting cell.^28^ In family F4, combination of *SCUBE2/BOC* variants was associated with additional variants in *SHH, STK36* (see below) and *WNT4*, a member of Wnt pathway, implicated in regulation of SHH signalling^19^. In family F22, the *SCUBE2* variant results in a premature stop codon at position 525 (*Figure S7*), which results in truncation of its CUB domain and is predicted to directly affect its SHH-related activity.^28^ This family presented an additional candidate variant in *HIC1*, which genetically interacts with *PTCH1*.^29^ Mice deficient for *HIC1* exhibit craniofacial defects including HPE.^30^

The reported variant combinations were observed exclusively in the affected probands and were absent in asymptomatic individuals. Altogether, these results reveal recurrent mutations in *SCUBE2/BOC* and further strengthen the oligogenic inheritance model of HPE.

### Implication of primary cilium in HPE

Remarkably, 5 families presented candidate variants in genes related to the primary cilium - *STK36, IFT172, B9D1, MKS1, TCTN3* and *TULP3* (Figure 2C). Ciliary proteins are known to play essential roles in the transduction of SHH signalling downstream of PTCH1 during forebrain development.^19,21^

*STK36*, also known as *Fused*, is a ciliary protein implicated in SHH signalling and associated to craniofacial phenotypes.^19,21^ *IFT172* codes for a core component of intraflagellar transport complex IFT-B required for ciliogenesis and regulation of SHH signal transduction. Moreover, *IFT172^−/−^* mice exhibit reduced expression of *Shh* in the ventral forebrain and severe craniofacial malformations including HPE.^20^ *B9D1, MKS1* and *TCTN3* are all members the transition zone protein complex implicated in regulation of ciliogenesis.^31^ The disruption of *B9D1* and *MKS1* in mouse models causes craniofacial defects that include HPE.^32,33^ Although no mouse model is available for *TCTN3*, its expression profile is highly similar to that of *SHH* and disruption of its protein complex partners (*TCTN1*, *TCTN2, CC2D2A, MKS1, B9D1*) leads to HPE in mouse.^31–33^ Moreover, *TCTN3* was shown to be necessary for the transduction of SHH signal and *TCTN3* mutations were found in patients affected by ciliopathies.^34^ Finally, *TULP3* is a critical repressor of *Shh* signalling in mouse and is associated to various craniofacial defects.^19^

Additional variants observed in these families include a heterozygous deletion of *SIX3*, missense mutations in *SHH, SCUBE2, BOC* and *LRP2* (described above) as well as two genes implicated in PCP pathway (Figure 3)*: CELSR1* (2 families) and *PRICKLE1*, both associated with craniofacial defects in mouse mutants (Figure 2C).^19,21,35^ Similarly to previously described cases, the oligogenic events were present exclusively in the affected probands.

Given the essential role of the primary cilium in SHH signal transduction, these observations strongly suggest that rare variants in ciliary genes contribute to the disease onset in these families.

### Correlation between variant combinations and secondary clinical features

To provide additional evidence, we performed an in-depth analysis of secondary clinical features associated with HPE in our patients. Deep clinical phenotyping identified clinical similarities between unrelated patients (Table 2) as well as overlaps of secondary clinical features between patients and the corresponding mouse mutants. Noteworthy, the two unrelated patients having variants in *FAT1* and *NDST1* shared a large set of specific secondary clinical features, including mandibular and ear abnormalities. Intrauterine growth restriction (IUGR) was found exclusively in the two patients with *COL2A1* variants. The most severely affected child in family F16 (*FAT1/NDST1/COL2A1*) presented a strong overlap with *NDST1-null* and *COL2A1*-null mutant mice (HPE, mandibular anomalies, absent olfactory bulb, abnormal nose morphology).^24,25^ Similarly, proboscis and eye defects were observed in both *FAT1/NDST1/SHH* patient and *FAT1^−/−^* mice.^22^

The 2 unrelated *SCUBE2/BOC* cases in families F4 and F22 presented cebocephaly, a midline facial anomaly characterized by ocular hypotelorism and a single nostril, which was absent in all other patients. Consistently, *SCUBE2* is highly expressed in the nasal septum in mouse,^36^ and cebocephaly was previously associated with *CDON –* another known HPE gene sharing highly similar functions and structure with *BOC.^37^*

Finally, the 2 patients with variants in ciliary genes (*IFT172/PRICKLE1* and *SIX3/TCTN3/TULP3*) both presented polydactyly, a clinical feature commonly associated with ciliopathies.^21^ Importantly, the patient with the oligogenic combination *IFT172/PRICKLE1* presented a large set of overlapping clinical features with the corresponding mouse mutants including polydactyly, cleft palate and eye defects, suggesting that these variants contribute to disease onset in this patient.^20,35^

While these clinical features are not specific to HPE, the described overlaps provide additional support for disease implication of the presented candidate variants.

## Discussion

In this study, we addressed the relevance of oligogenic model for unsolved HPE cases. We provide evidence that the onset of HPE arises from the combined effects of hypomorphic variants in several genes belonging to critical biological pathways of brain development. To circumvent the limitations of classical WES analysis in complex rare disorders, we combined clinically-driven and co-expression network analyses with classical WES variant prioritization. This strategy was applied to 26 HPE families and allowed prioritization of 180 genes directly linked to the SHH signalling, cilium and Wnt/PCP pathways (Figure 3). The analysis of oligogenic events in patients with HPE anomalies revealed 19 genes including 15 genes previously unreported in human HPE patients (Table 2). All these genes are either associated with HPE phenotypes in corresponding mouse models (such as *FAT1, NDST1*), present highly similar expression patterns with already known HPE genes in the developing brain (such as *SCUBE2, TCTN3*) or both. We observed co-occurrence of mutations in several gene pairs such as *FAT1/NDST1* and *SCUBE2/BOC*, which provides additional arguments towards their implication in HPE. We additionally show that in-depth evaluation of secondary clinical features in patients with HPE anomalies and comparison to published mouse knockout models allow the identification of recurrent *genotype-phenotype* correlations that provide additional arguments for their pathogenicity.

The main challenge in disease-gene discovery by Whole Exome Sequencing is to identify disease-related variants among a large background of non-pathogenic polymorphisms.^8,18^ For example, the presented *FAT1* encodes a large protocadherin gene spanning over 139 kb in the human genome and presenting over 2000 missense variants with MAF below 1 % in the gnomAD database. Despite this high number of variations found in the general population, rare variants in *FAT1* were recently shown to be causative in several genetic disorders including facioscapulohumeral dystrophy-like disease.^38^ Hence, correct interpretations and conclusions require extremely careful assessment of available biological and clinical knowledge.

To improve the pertinence of our study, we developed a strategy to restrict the potential candidates by targeting genes with biological and clinical arguments for their implication in the disease. Implication of a given gene in a disease is often supported by the similarity between the human pathology and the phenotype obtained in relevant animal models.^18^ Accordingly, in this study, the main evidence of causality for candidate genes was that their disruption leads to clinically-defined HPE-related phenotypes in corresponding published mutant mouse models. Unlike other phenotypes such as reduced body weight, holoprosencephaly is a rare effect of gene knockout in mice as it is present in less than 1 % of genes (as reported in the MGI database). Recent exome sequencing studies have applied similar phenotype-driven approaches to identify causal variants in monogenic disorders. Dedicated tools have been developed in that aim (Exomiser, Phive)^10^ but none are designed for non-Mendelian traits involving hypomorphic variants with mild effects. We provide a method to specifically address such cases and show that further developments are necessary to improve the diagnosis of genetic disorders especially by taking into account oligogenic inheritance. Inclusion of carefully defined mouse mutant phenotypes is of powerful value as certain phenotypes like HPE are very informative due to their rarity.

Prioritization tools can also include protein–protein interaction (PPI) networks information, which improves performance in cases where candidate genes do not have an associated knockout mouse model. However, PPI-based prioritization is limited when disease investigation requires incorporation of tissue-specific data. The key process affected by HPE is the elaboration of the forebrain and its dorso-ventral patterning.^40^ Deciphering the biological mechanisms involved in the early brain development is therefore necessary to provide relevant information to select disease-related genes. To incorporate tissue-specificity, we performed analysis using the RNA-Seq data of embryonic human brain at the earliest available developmental stages (from 4 to 17 pcw) as provided by the Human Development Biology Resource.^17^ We defined relevant co-expression modules and selected candidate genes of which expression patterns follow those of known HPE genes. Further analysis showed that the resulting candidate genes, such as *SCUBE2* and *TCTN3*, are pertinent as they are equally implicated in the SHH pathway that is the primary HPE pathway.^28,34^ Co-expression analysis provides additional insight into disease pathogenesis by establishing the first link between previously unrelated genes. A future challenge will be to generalise this approach but such a task will face the necessity to incorporate disease relevant co-expression modules that need to be pre-computed.

Patients exhibiting HPE-anomalies present enrichment of rare variants in genes related to the SHH pathway, as well as to the Wnt/PCP and primary cilia pathways, which were both shown to functionally interact with and regulate SHH pathway.^19–21,33^ Accumulation of multiple rare variants in genes related to these pathways will likely disrupt the dorso-ventral gradient of the SHH morphogen^40^, leading to an incomplete cleavage of the forebrain and, ultimately, to HPE. In this model, distinction between different manifestations of HPE lies in the degree of overall functional impact on SHH signalling.^7^ Moreover, depending on the affected genes and pathways, HPE patients would present different secondary clinical features.

The observed overlapping secondary clinical features further support the causality of the reported variants for HPE. As hypomorphic mutations do not have the same impact as the complete inactivation of a gene in most cases, phenotypic overlaps may be challenging to detect and require expert assessment of clinical and biological data. For example, mice deficient in *NDST1* exhibit agnathia^24^ (absence of the lower jaw) while unrelated patients presenting candidate variants in *NDST1* exhibit respectively prognathia and retrognathia (abnormal positioning of the lower jaw). All three phenotypes are part of the same spectrum of mandibular anomalies. From a clinical perspective, overlap of secondary clinical features between the patient and the animal models provides additional critical evidence of a causal relationship between candidate gene and disease. Key issue here remains the semantic representation of patient’s phenotype and the use of a well-established phenotypic ontology during the examination processes. Explorations of secondary clinical features should be performed in future studies of genetic diseases.

Additional molecular screenings in larger populations of HPE patients are necessary to definitely assess the implication of our candidate genes in the disease. Therefore, we propose to include these novel genes into future genetic screenings of HPE patients.

In conclusion, this paper presents novel genes implicated in HPE and illustrates that HPE presents an oligogenic inheritance pattern requiring the joint effect of multiple genetic variants acting as hypomorphic mutations. The proposed inheritance pattern accounts for a wide clinical spectrum of HPE and explains the significant part of cases in which no molecular diagnosis could be established by conventional approaches. It also explains the incomplete penetrance and variable expressivity of inherited causal mutations observed in the reported cases of HPE.^1,4^ We propose that in cases of non-Mendelian diseases with variable phenotypes, the possibility of oligogenic inheritance needs to be evaluated. Exploration of such events will improve the diagnostic yield of complex developmental disorders and will contribute to better understand the mechanisms that coordinate normal and pathological embryonic development.

## Acknowledgements

This work was supported by La Fondation Maladie Rares and the Agence de la Biomedecine. The authors acknowledge the Centre de Ressources Biologiques (CRB)-Santé (http://www.crbsante-rennes.com) of Rennes for managing patient samples. We would like to thank the families for their participation in the study, all clinicians who referred HPE cases, the eight CLAD (Centres Labellisés pour les Anomalies du Développement) within France that belong to FECLAD, French centers of prenatal diagnosis (CPDPN) and the SOFFOET for fetus cases, and the “filière AnDDI-Rares.” We particularly thank all members of the Molecular Genetics Laboratory (CHU, Rennes) and of the Department of Genetics and Development (UMR6290 CNRS, Université Rennes 1) for their help and advice.

## References

1. Solomon BD, Gropman A, Muenke M. Holoprosencephaly Overview. In: Adam MP, Ardinger HH, Pagon RA, et al., eds. GeneReviews^®^. Seattle (WA): University of Washington, Seattle; 1993. Last update: August 29, 2013

2. Mouden C, Tayrac M de, Dubourg C, et al. Homozygous STIL Mutation Causes Holoprosencephaly and Microcephaly in Two Siblings. PLOS ONE. 2015;10(2):e0117418.

3. Mouden C, Dubourg C, Carré W, et al. Complex mode of inheritance in holoprosencephaly revealed by whole exome sequencing. Clin Genet. 2016;89(6):659–668.

4. Mercier S, Dubourg C, Garcelon N, et al. New findings for phenotype-genotype correlations in a large European series of holoprosencephaly cases. J Med Genet. 2011;48(11):752–760.

5. Dubourg C, Carré W, Hamdi-Rozé H, et al. Mutational Spectrum in Holoprosencephaly Shows That FGF is a New Major Signaling Pathway. Hum Mutat. 2016;37(12):1329–1339.

6. Dupé V, Rochard L, Mercier S, et al. NOTCH, a new signaling pathway implicated in holoprosencephaly. Hum Mol Genet. 2011;20(6):1122–1131.

7. Mercier S, David V, Ratié L, Gicquel I, Odent S, Dupé V. NODAL and SHH dose-dependent double inhibition promotes an HPE-like phenotype in chick embryos. Dis Model Mech. 2013;6(2):537–543.

8. Bamshad MJ, Ng SB, Bigham AW, et al. Exome sequencing as a tool for Mendelian disease gene discovery. Nat Rev Genet. 2011;12(11):745–755.

9. Philippakis AA, Azzariti DR, Beltran S, et al. The Matchmaker Exchange: a platform for rare disease gene discovery. Hum Mutat. 2015;36(10):915–921.

10. Smedley D, Jacobsen JOB, Jäger M, et al. Next-generation diagnostics and disease-gene discovery with the Exomiser. Nat Protoc. 2015;10(12):2004–2015.

11. Li L, Bainbridge MN, Tan Y, Willerson JT, Marian AJ. A Potential Oligogenic Etiology of Hypertrophic Cardiomyopathy: A Classic Single-Gene Disorder. Circ Res. 2017;120(7):1084–1090.

12. Stark Z, Dashnow H, Lunke S, et al. A clinically driven variant prioritization framework outperforms purely computational approaches for the diagnostic analysis of singleton WES data. Eur J Hum Genet EJHG. 2017;25(11):1268–1272.

13. Lee H, Deignan JL, Dorrani N, et al. Clinical exome sequencing for genetic identification of rare Mendelian disorders. JAMA. 2014;312(18):1880–1887.

14. Richards S, Aziz N, Bale S, et al. Standards and guidelines for the interpretation of sequence variants: a joint consensus recommendation of the American College of Medical Genetics and Genomics and the Association for Molecular Pathology. Genet Med Off J Am Coll Med Genet. 2015;17(5):405–424.

15. Smith CL, Blake JA, Kadin JA, Richardson JE, Bult CJ, Mouse Genome Database Group. Mouse Genome Database (MGD)-2018: knowledgebase for the laboratory mouse. Nucleic Acids Res. 2018;46(D1):D836–D842.

16. Langfelder P, Horvath S. WGCNA: an R package for weighted correlation network analysis. BMC Bioinformatics. 2008;9:559.

17. Lindsay SJ, Xu Y, Lisgo SN, et al. HDBR Expression: A Unique Resource for Global and Individual Gene Expression Studies during Early Human Brain Development. Front Neuroanat. 2016; 10.

18. MacArthur DG, Manolio TA, Dimmock DP, et al. Guidelines for investigating causality of sequence variants in human disease. Nature. 2014;508(7497):nature13127.

19. Murdoch JN, Copp AJ. The relationship between Sonic hedgehog signalling, cilia and neural tube defects. Birt Defects Res A Clin Mol Teratol. 2010;88(8):633–652.

20. Gorivodsky M, Mukhopadhyay M, Wilsch-Braeuninger M, et al. Intraflagellar Transport Protein 172 is essential for primary cilia formation and plays a vital role in patterning the mammalian brain. Dev Biol. 2009;325(1):24–32.

21. Goetz SC, Ocbina PJR, Anderson KV. The Primary Cilium as a Hedgehog Signal Transduction Machine. Methods Cell Biol. 2009;94:199–222.

22. Ciani L, Patel A, Allen ND, ffrench-Constant C. Mice lacking the giant protocadherin mFAT1 exhibit renal slit junction abnormalities and a partially penetrant cyclopia and anophthalmia phenotype. Mol Cell Biol. 2003;23(10):3575–3582.

23. Rock R, Schrauth S, Gessler M. Expression of mouse dchs1, fjx1, and fat-j suggests conservation of the planar cell polarity pathway identified in drosophila. Dev Dyn. 2005;234(3):747–755.

24. Grobe K. Cerebral hypoplasia and craniofacial defects in mice lacking heparan sulfate Ndst1 gene function. Development. 2005;132(16):3777–3786.

25. Leung AWL, Wong SYY, Chan D, Tam PPL, Cheah KSE. Loss of procollagen IIA from the anterior mesendoderm disrupts the development of mouse embryonic forebrain. Dev Dyn. 2010;239(9):2319–2329.

26. Christ A, Christa A, Kur E, et al. LRP2 is an auxiliary SHH receptor required to condition the forebrain ventral midline for inductive signals. Dev Cell. 2012;22(2):268–278.

27. Hong M, Srivastava K, Kim S, et al. BOC is a modifier gene in holoprosencephaly. Hum Mutat. 2017;38(11):1464–1470.

28. Jakobs P, Exner S, Schürmann S, et al. Scube2 enhances proteolytic Shh processing from the surface of Shh-producing cells. J Cell Sci. 2014;127(Pt 8):1726–1737.

29. Briggs KJ, Corcoran-Schwartz IM, Zhang W, et al. Cooperation between the Hic1 and Ptch1 tumor suppressors in medulloblastoma. Genes Dev. 2008;22(6):770–785.

30. Carter MG. Mice deficient in the candidate tumor suppressor gene Hic1 exhibit developmental defects of structures affected in the Miller-Dieker syndrome. Hum Mol Genet. 2000;9(3):413–419.

31. Garcia-Gonzalo FR, Corbit KC, Sirerol-Piquer MS, et al. A transition zone complex regulates mammalian ciliogenesis and ciliary membrane composition. Nat Genet. 2011;43(8):776–784.

32. Dowdle WE, Robinson JF, Kneist A, et al. Disruption of a Ciliary B9 Protein Complex Causes Meckel Syndrome. Am J Hum Genet. 2011;89(1):94–110.

33. Wheway G, Abdelhamed Z, Natarajan S, Toomes C, Inglehearn C, Johnson CA. Aberrant Wnt signalling and cellular over-proliferation in a novel mouse model of Meckel-Gruber syndrome. Dev Biol. 2013;377(1):55–66.

34. Thomas S, Legendre M, Saunier S, et al. TCTN3 mutations cause Mohr-Majewski syndrome. Am J Hum Genet. 2012;91(2):372–378.

35. Yang T, Jia Z, Bryant-Pike W, et al. Analysis of PRICKLE1 in human cleft palate and mouse development demonstrates rare and common variants involved in human malformations. Mol Genet Genomic Med. 2014;2(2):138–151.

36. Xavier GM, Cobourne MT. Scube2 expression extends beyond the central nervous system during mouse development. J Mol Histol. 2011;42(5):383–391.

37. Zhang W, Kang J-S, Cole F, Yi M-J, Krauss RS. Cdo functions at multiple points in the Sonic Hedgehog pathway, and Cdo-deficient mice accurately model human holoprosencephaly. Dev Cell. 2006;10(5):657–665.

38. Puppo F, Dionnet E, Gaillard M-C, et al. Identification of Variants in the 4q35 Gene FAT1 in Patients with a Facioscapulohumeral Dystrophy-Like Phenotype. Hum Mutat. 2015;36(4):443–453.

39. Reed DR, Lawler MP, Tordoff MG. Reduced body weight is a common effect of gene knockout in mice. BMC Genet. 2008;9:4.

40. Fernandes M, Hébert JM. The ups and downs of holoprosencephaly: dorsal versus ventral patterning forces. Clin Genet. 2008;73(5):413–423.

